# Ketone Bodies-mediated Cysteine Modifications Discovered by Chemical Proteomics

**DOI:** 10.1101/2025.05.25.655943

**Authors:** Yuan-Fei Zhou, Ling Zhang, Zhuoyi Niu, Xin Wang, Ryan Hunt, Yingming Zhao, Nima Sharifi, Zhipeng A. Wang

## Abstract

All the studies of ketone body-dependent post-translational modifications (PTMs), notably those mediated by ketone bodies, β-hydroxybutyrate (Bhb) and acetoacetate (Acac), have focused on lysine acylations. However, given the chemically diverse and reactive nature of metabolites generated, it remains unclear whether non-lysine modifications can also happen. Here, we report the synthesis of an acetoacetate-alkyne (Acac-alkyne) chemical probe that enables efficient metabolic labeling, robust fluorescent visualization, and mass spectrometry-based identification of Acac-modified proteins. By combining chemical proteomics with open-search strategy, we showed that Acac will induce previously uncharacterized cysteine modifications in mammalian cells. Notably, cysteine S-crotonation (Ccr) is validated by employing both probe-based and standard peptide-based co-elution assays. Metabolic pathway tracing further identifies BDH1 and ECHS1 as key enzymes that generate Ccr formation. Together, these findings establish ketone metabolism as a novel source of cysteine modifications and provide an alternative mechanistic pathway to explain the profound biological effects of ketone body.

## Introduction

Ketogenesis, triggered by fasting, exercise, or a ketogenic diet, increases circulating ketone bodies β-hydroxybutyrate (Bhb) and acetoacetate (Acac) to millimolar level. ^1, 2^ Ketone bodies are derived from acetyl-CoA and serve as metabolically efficient alternative energy sources, especially under conditions of glucose deprivation.^3^ After circulation, Bhb and Acac are taken up by extrahepatic tissues and converted into acetyl-CoA, which subsequently enters the tricarboxylic acid cycle to produce ATP in tissues.^4^ In addition to serving as metabolic fuels, these ketones are converted to CoA derivatives that can be installed on histones and non-histone proteins, leading to two recently identified marks: lysine β-hydroxybutyrylation (Kbhb) and lysine acetoacetylation (Kacac).^5, 6^ These discoveries demonstrate that ketone bodies directly reshape the epigenetic landscape, bridging systemic metabolic status with gene regulation.^7-9^ Despite these advances, the effects of ketone-driven PTMs beyond lysine acylation remain largely unexplored, contributing to the incomplete understanding of ketogenesis at the molecular level.

To address this gap, we combine the chemical proteomics with open-search strategy to investigate if ketone bodies can be precursors for protein modifications other than lysine acylation. To this end, we synthesized a bioorthogonal probe of Acac (Acac-alkyne) that closely mimics the chemical structure and biochemical reactivity of Acac. We then performed chemical proteomic profiling of Acac modified proteins in living cells in a site-specific manner, unveiling previous unknown cysteine modifications (Cys427, Cys267) induced by ketone body via open-search. In order to confirm the modification structure of Cys267, we used a probe-based co-elution assay to confirm that Acac transfer into crotonyl intermediate to form a novel cysteine S-crotonation (Ccr). To demonstrate that this modification exists endogenously, we further confirmed this modification by PRM scanning and co-elution experiments with synthetic peptides. We next sought to determine how Ccr is formed endogenously within cells. We proposed that Ccr is generated though Bhb and crotonyl intermediate. A probe-based co-elution assay confirmed that Bhb and crotonate will also form the same Ccr modifications. Knockout of two key enzymes, BDH1 and ECHS1, resulted in decreased in-gel fluorescence signals, indicating Ccr undergoes reverse β-oxidation to form Ccr. Collectively, our findings reveal ketone body as a previously unrecognized driver of new cysteine modifications and provide an alternative mechanistic pathway to explain the profound biological effects of ketone body.

## Results

Bioorthogonal chemical probes provide a robust platform for interrogating post-translational modifications (PTMs).^10, 11^ Probes bearing alkyne or azide tags can be metabolically incorporated into the corresponding native PTM sites on endogenous proteins in cells.^12-15^ A subsequent click reaction^16, 17^ conjugates fluorescent dyes or biotin to these tags, enabling selective visualization and identification of the labeled proteins. Ketogenesis, triggered by fasting, exercise, or a ketogenic diet, elevates circulating ketone body levels to the millimolar range through acetyl-CoA metabolism (Fig 1A). Accordingly, we hypothesize that an alkynyl analogue of Acac will enter this metabolic pathway and incorporate into Acac-modified proteins in live cells. We thus designed and synthesized Acac derivative probe, Acac-alkyne, containing a terminal alkynyl group (Fig 1B). Acac-alkyne was synthesized from 4-pentynoic acid (compound **1**) and Meldrum’s acid to generate compound **2** (Fig S1A), which was converted to compound **3** through *tert-*butanol mediated esterification, and subsequent deprotection to form the final Acac-alkyne probe (compound **4**).

**Fig 1.**
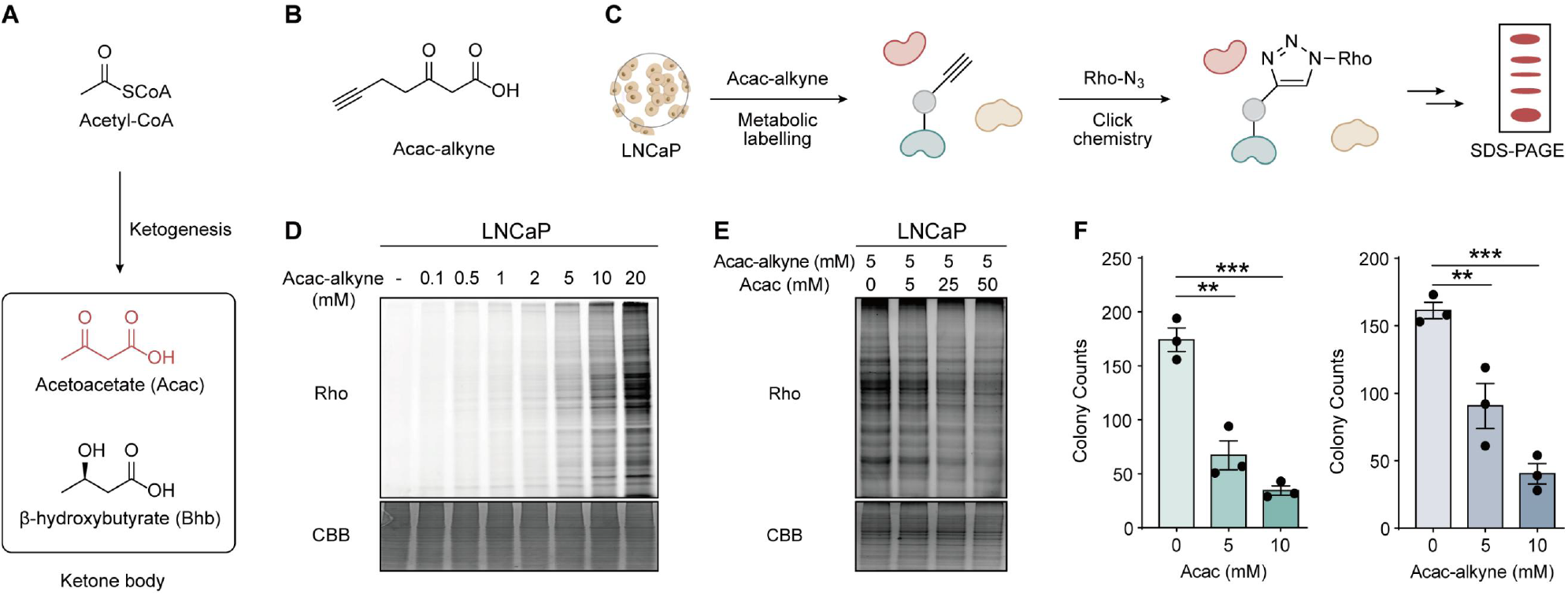
Design, synthesis and in-gel fluorescence of Acac-alkyne probe for profiling Acac modified proteins. (A) The ketone body produced from acetyl-CoA via ketogenesis metabolism. (B) Design of the bioorthogonal Acac derivative probe, Acac-alkyne. (C) In-gel fluorescence strategy for detection of Acac-modified proteins using Acac-alkyne. (D) LNCaP cells were labeled with increasing concentrations of Acac-alkyne, showing a dose-dependent manner. (E) Co-treatment with increasing concentrations of Acac attenuated the labeling signal of Acac-alkyne. (F) Acac and Acac-alkyne exhibited similar colony-suppressive effects.

With Acac-alkyne in hand, we first evaluated whether this probe can be metabolically incorporated into proteins in mammalian cells. To this end, LNCaP prostate cancer cells were treated with Acac-alkyne for 24 hours. After harvesting the cells, the whole cell lysates were subjected to azide-alkyne click chemistry to conjugate the Acac-alkyne labeled proteins to a rhodamine dye. The labeled proteins were then resolved by SDS-PAGE gel and visualized by in-gel fluorescence (Fig 1C). Our experiment demonstrated that a wide range of proteins was labeled by Acac-alkyne in a dose-dependent manner, with 5-10 mM as optimal concentrations (Fig 1D). Similar metabolic labeling results were observed across multiple cell types, including HEK293T, 22Rv1, Hepa1-6 and NIH3T3 (Fig S1B). Our result highlights the general applicability of Acac-alkyne and excellent labeling efficiency. In addition, when cells were treated with a mixture of Acac and Acac-alkyne for 2 hours, the fluorescence signal of the probe-labeled proteins was decreased with increasing concentration of Acac in a dose-dependent manner. This result indicated that Acac-alkyne is metabolized through the native Acac metabolic pathway and incorporated into Acac-modified proteins (Fig 1E). Functionally, Acac has been reported to significantly reduce colony formation in cancer cells.^18^ Consistently, we also observed that Acac significantly inhibited colony formation after treatment, and Acac-alkyne recapitulated this effect (Fig 1F). In contrast, Bhb showed no such activity (Fig S1C, D), indicating that Acac-alkyne mimics Acac, but not Bhb. Collectively, these findings validated the Acac-alkyne as a bioorthogonal chemical probe to profile Acac-modified proteins.

After verifying the effectiveness of Acac-alkyne probe, we performed site-specific chemical proteomics study to unveil Acac-modified proteins at the proteome level by tandem orthogonal proteolysis–activity-based protein profiling (TOP-ABPP) methods.^19, 20^ Lysates from Acac-alkyne-labelled LNCaP cells were prepared, conjugated with the acid-cleavable azido-DADPS-biotin tag, followed by affinity purification, on-bead digestion and acid-mediated cleavage to enrich modified peptides (Fig 2A). The eluted Acac-alkyne modified peptides were analyzed by LC-MS/MS. We first performed an open-search strategy^21, 22^ to identify any potential modifications that may occur. As anticipated, we have found lysine-derived modification representing Kacac (K, +265.1426 Da) (Fig 2B). Notably, we also found two previously unknown cysteine modifications with adduct masses are cysteine plus +267.1577 Da (Cys267) and +427.1867 Da (Cys427) (Fig 2B). These results drew our attention, as Acac has not previously been reported to induce any cysteine modifications.

**Fig 2.**
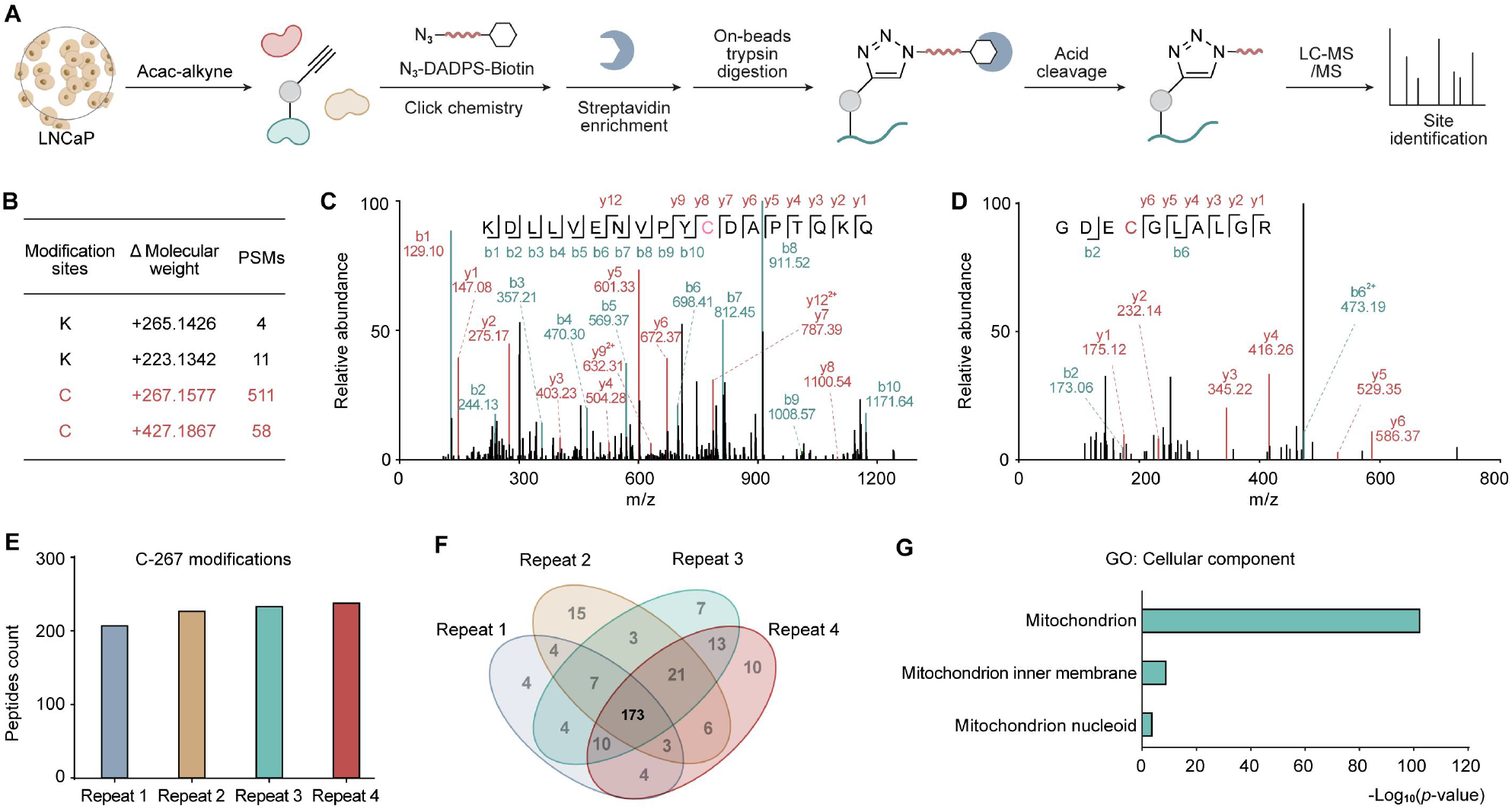
Combination chemical proteomic and open-search strategy to characterize unknown cysteine modifications. (A) Workflow for the site-specific chemical profiling of Acac-modified proteins using Acac-alkyne. (B) Open-search strategy characterized two unknown cysteine modifications. (C, D) A representative MS2 spectrum shows the Cys267 (LYRM7_C96) and Cys427 (ALDH18A1_C247) modifications, respectively. (E, F) The chemical proteomics data indicated 173 peptides containing Cys267 modification across four replicates. (G) GO enrichment analysis of identified Cys267 modifications are more represented in the mitochondrion.

We manually verified the representative MS2 spectra for the modified peptides, such as Cys267 modification at LYRM7_C96 (KDLLVENVPY**C**DAPTQKQ) and Cys427 modification at ALDH18A1_C247 (GDE**C**GLALGR), confirmed high confidence in identification (Fig 2C, D). To improve reliability and coverage of the modified cysteine identification, we subsequently conducted a closed search with only the sites identified in all four biological replicates considered for further analysis. The refinement resulted in 173 peptides containing Cys267 modification (Fig 2E, F), corresponding to 120 distinct proteins (Fig S2A). Similarly, 35 peptides containing Cys427 modification, also mapping to 29 proteins were detected in all four biological repeats (Fig S2C, D). Cellular component analysis using the Gene Ontology (GO) database revealed that both Cys267 and Cys427 modifications are exclusively represented in the mitochondria (Fig 2G, Fig S2E), with associated biological processes involving tRNA processing and protein biosynthesis. Molecular function analysis the modified proteins enriched in ribosomal protein, ribonucleoprotein and oxidoreductase (Fig S2B, E). Collectively, combining chemical proteomic and open-search strategy, we characterized two novel cysteine modifications that exclusively distributed in mitochondria, significantly expanding the chemical space of ketone body-derived PTMs and its potential role in metabolic regulation.

Given that Cys267 exhibited the highest number of modification sites, we prioritized its structural characterization. Based on the observed Cys267 mass shift, there are two potential candidates (Fig 3A) formed via different metabolic pathways. For candidate 1, Acac-alkyne was hypothesized to undergo carbonyl reduction followed by thioester formation with protein cysteine residues to generate Cys267 modification (Fig S3A). However, thioesters are generally unstable under basic conditions and prone to hydrolysis. To assess this possibility, we treated Acac-alkyne labeled samples with or without 200 mM hydrazine^23^ (Fig S3B), and observed no change in fluorescence intensity, thereby excluding a thioester intermediate.

**Fig 3.**
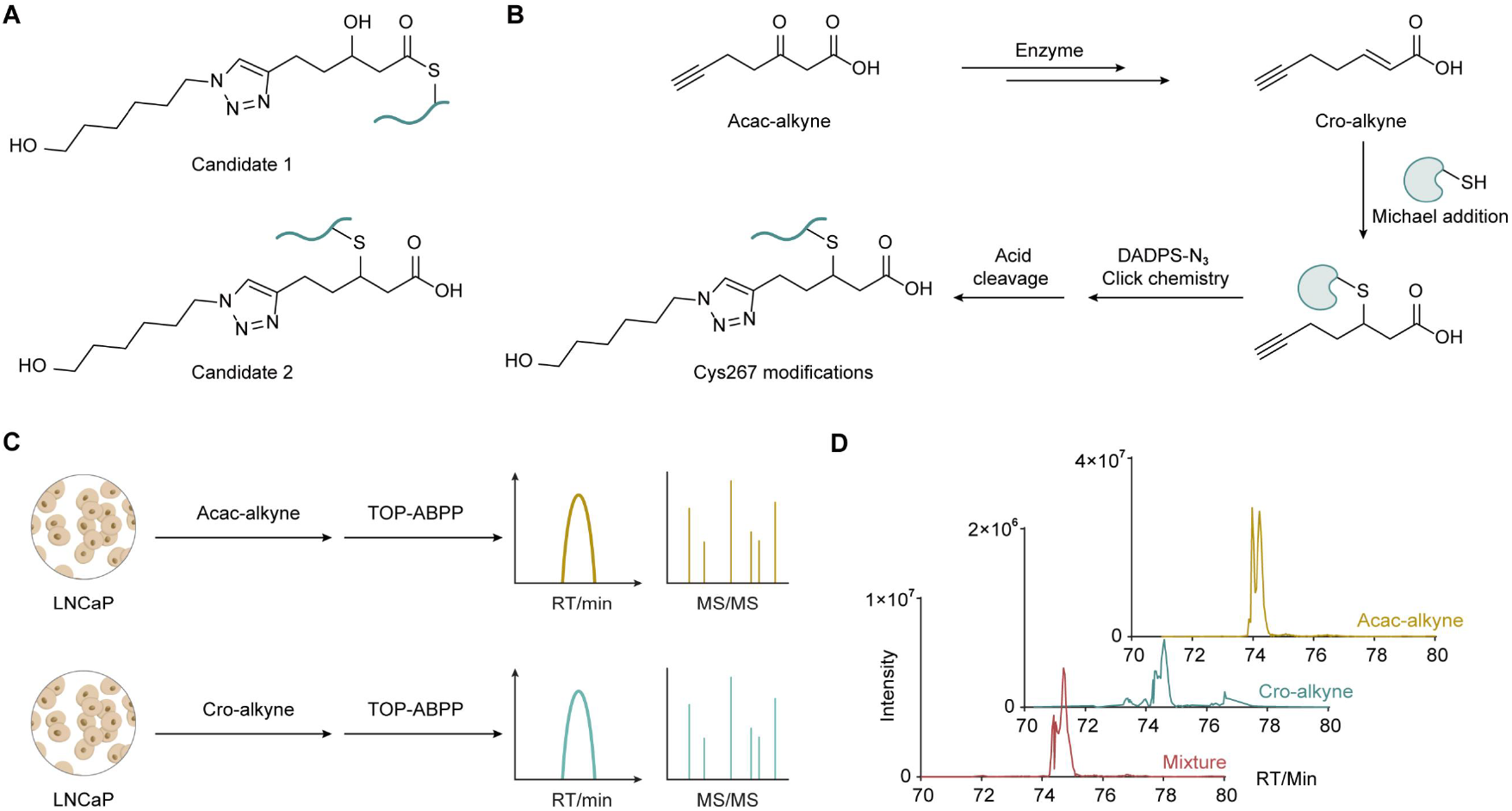
Confirmation the structure of Cys267 modification by probe-based co-elution assay. (A) Structures of two potential candidates bearing Cys267 modification. (B) Acac-alkyne proposed transformed to Cro-alkyne, followed by Michael addition with cysteine residues to form Cys267 modification. (C) Comparing retention time and MS characteristics of Cro-alkyne and Acac-alkyne modified peptides by TOP-ABPP assay. (D) Retention time of the extracted ion chromatogram corresponding to the Cys267 modification of PRDX3_C229 in Acac-alkyne, Cro-alkyne and mixture groups.

We next evaluated candidate 2, in which Acac-alkyne may undergo transformation to Cro-alkyne, followed by Michael addition to cysteine residues (Fig 3B). If it is the case, we postulated that the corresponding alkyne-containing mimic of crotonate (Cro-alkyne)^15^ could also form the same Cys267 modification. To test this, we synthesized Cro-alkyne and performed the TOP-ABPP assay to compare its labeling profile with that of Acac-alkyne (Fig 3C). Among 133 peptides modified by Cro-alkyne at Cys267, 24 sites exhibited co-elution and matched MS2 fragmentation patterns with peptides labeled by Acac-alkyne, indicating shared modification sites and suggesting overlapping reactivity profiles between the two probes at these positions (Fig S3C). Taking PRDX3_C229 as an example, the modified peptide (AFQYVETHGEV**C**PANWTPDSPTIKPSPAASK) showed the same retention times (Fig 3C) and fragmentation patterns in Acac-alkyne and Cro-alkyne groups (Fig S3D). Collectively, these results support candidate 2 as the novel cysteine modification structure, where Acac-alkyne is converted to Cro-alkyne, leading to Cys267 modification through Michael addition.

After confirming the conversion of Acac-alkyne to Cro-alkyne, we hypothesized that endogenous Acac may also undergo a similar transformation to generate crotonate intermediate, resulting in a +86.0368 Da mass shift to cysteine residue (Fig 4A). We designated this modification as cysteine crotonation (Ccr). To test whether Acac treatment could induce Ccr under physiological conditions, we treated the cells with 10mM Acac for 24 hours, isolated mitochondria and prepared samples for targeted detection using a PRM inclusion list identified by Acac-alkyne metabolic labeling (Fig S4A). As a result, we detected three Ccr-modified peptides derived from ACAT1_C126, HSD17B10_C112, and PRDX3_C229 (Fig 4B).

**Fig 4.**
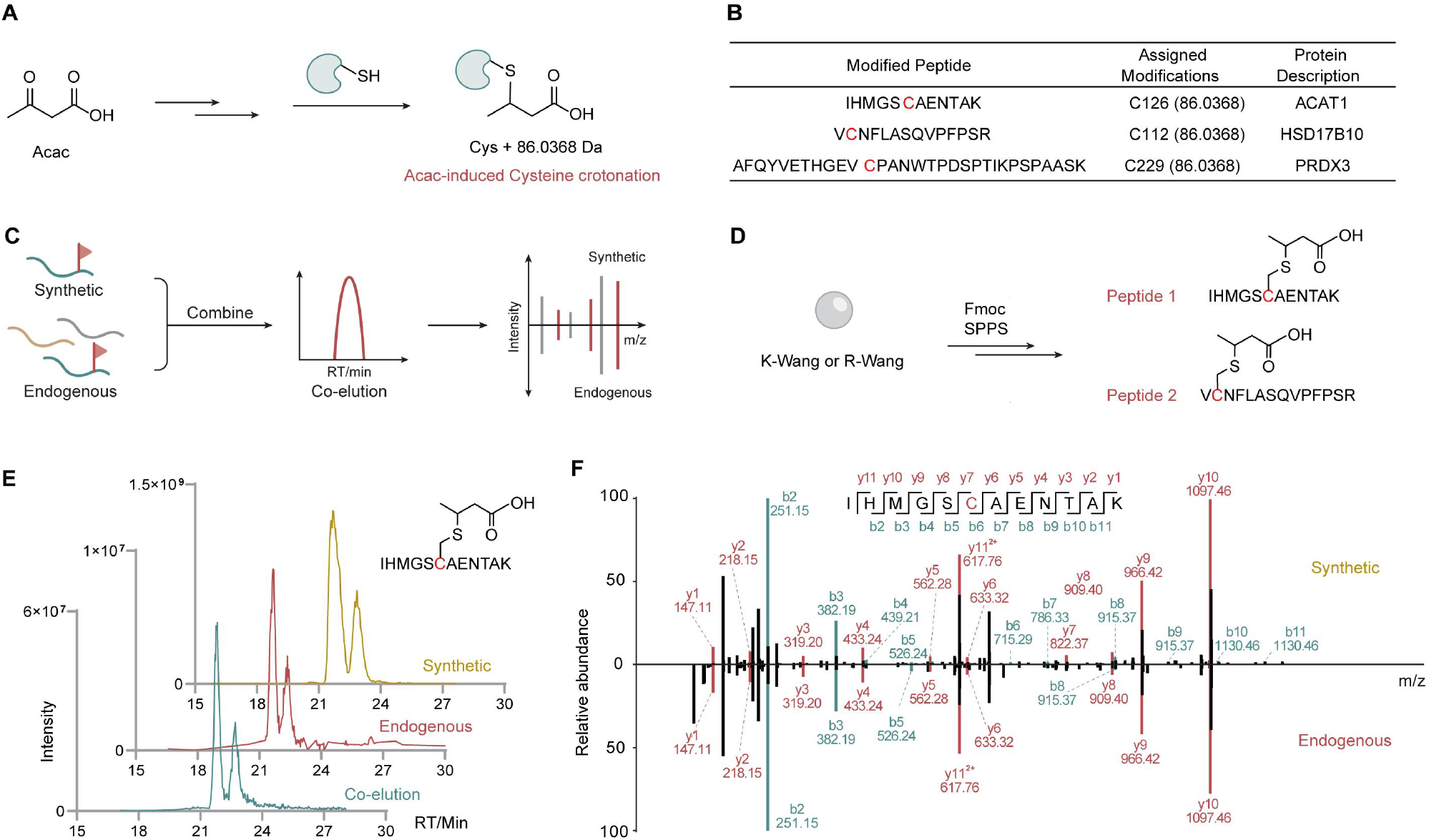
Identification the endogenous cysteine modification by standard peptides co-elution assay. (A) Acac is proposed to induce cysteine crotonation (Ccr). (B) Ccr-modified peptides derived from ACAT1_C126, HSD17B10_C112, and PRDX3_C229 were idenfied by PRM scanning. (C) To validate the authenticity of the modified peptides by co-elution assay. (D) Synthesis two modified peptides IHMGS**Ccr**AENTAK and V**Ccr**NFLASQVPFPSR by SPPS. (E) The extracted ion chromatogram corresponding to Ccr modification of ACAT1_C126 (IHMGS**Ccr**AENTAK). (F) MS2 spectra of the Ccr modification at ACAT1_C126 (IHMGS**Ccr**AENTAK).

To ultimately validate the authenticity of the modified peptides (Fig 4C), we performed the co-elution experiment with synthetic peptides and Acac-treated mitochondrial samples, which is widely used as the golden standard to verify the structure of new modifications.^24, 25^ We selected to synthesize two modified peptides IHMGS**Ccr**AENTAK and V**Ccr**NFLASQVPFPSR via solid-phase peptide synthesis (SPPS) and conducted co-elution experiments to confirm the identity of the endogenous peptides (Fig 4D). Briefly, K-Wang or R-Wang resin was coupled with C terminal amino acids and then reacted with Fmoc-Cys(S-Tmp)-OH.^26^ After Tmp de-protection by DTT, the resin was treated with *tert*-butyl crotonate and then subjected to conventional SPPS to complete the whole sequence with Ccr modification. The co-elution assay with mitochondrial samples and synthetic peptides showed that the synthetic standard IHMGS**Ccr**AENTAK (Fig 4E, F) and V**Ccr**NFLASQVPFPSR (Fig S4C, D) have the same retention time and MS2 fragmentation patterns as the endogenously Ccr-modified peptides. Notably, the IHMGS**Ccr**AENTAK peptide consistently revealed the same ion peaks but two distinct retention time, suggesting that the Ccr modification exists as both *R*- and *S*-stereoisomers. This observation supports that the Michael addition step involved in Ccr formation proceeds without stereoselectivity. Collectively, these results confirm that endogenous Ccr modification exists in cells in response to Acac treatment.

To investigate the endogenous origin of Ccr, we next examined the potential biochemical pathway leading to its formation. Chemically, Acac is expected to be reduced to Bhb first, which may subsequently undergo dehydration to generate the crotonyl intermediate (Fig 5A). This raises a key mechanistic question: if this pathway is operative, could all Acac, Bhb and crotonate serve as precursors for Ccr? To test this hypothesis, we demonstrated that both Cro-alkyne and crotonate can undergo spontaneous, non-enzymatic Michael addition reactions with GSH (Fig S5A, B), indicating non-enzymatic step between crotonyl intermediate with cysteine residue. To next test whether Bhb can induce Ccr formation, we synthesized an alkyne-containing Bhb derivative (Bhb-alkyne) probe (Fig S5C) and performed a series of co-elution experiments using all three probes. These experiments confirmed that Acac-, Bhb-, and crotonate-derived probes can modify the same cysteine sites at ACAT1_C126, PRDX3_C229 and HSD17B10_C112 (Fig 5B, Fig S5D, E) and share reliable MS2 fragmentation spectra (Fig S5C), supporting Bhb as a viable Ccr precursor and showing the plausibility of a shared metabolic pathway with Acac.

**Fig 5.**
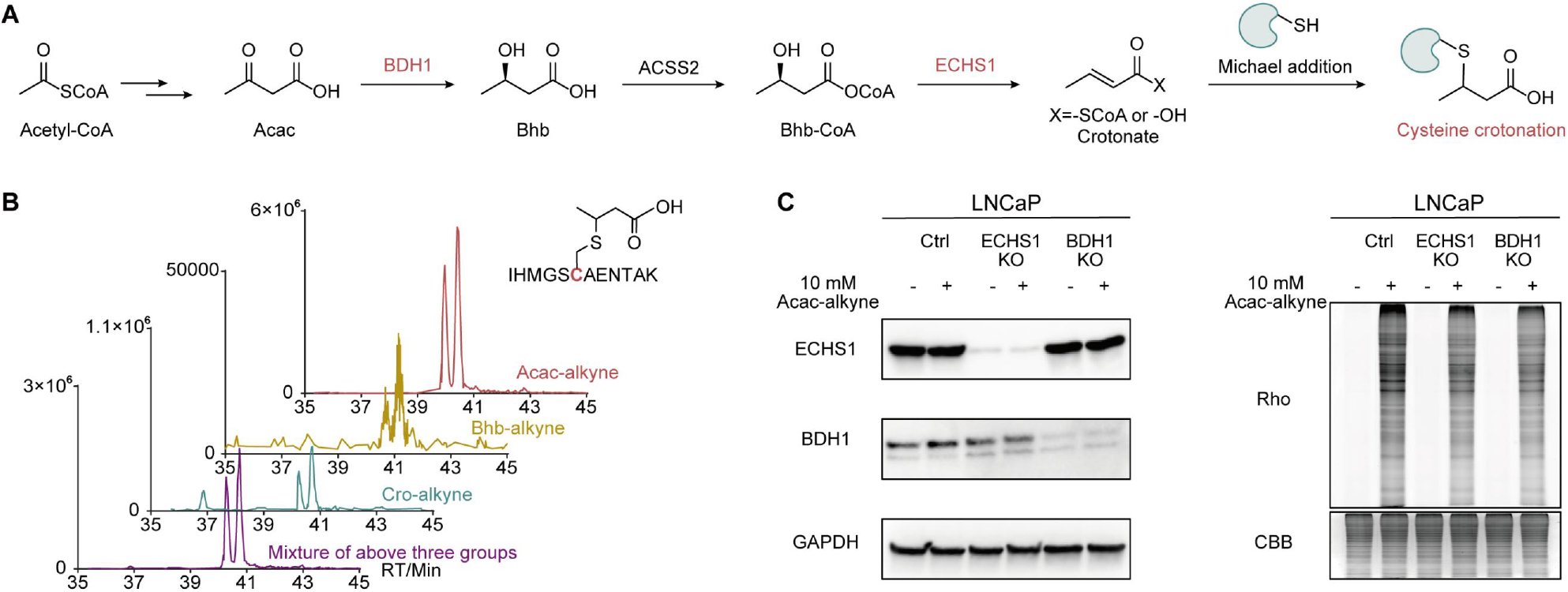
Exploration the metabolic pathway for Ccr formation. (A) Acac is expected to be first reduced to Bhb, which may subsequently undergo dehydration to generate crotonyl intermediate, then induce cysteine crotonation (Ccr). (B) The extracted ion chromatogram corresponding to ACAT1_C126 in Acac-alkyne, Bhb-alkyne, Cro-alkyne and mixture groups. (C) BDH1 and ECHS1 KO cell lines exhibited reduced fluorescence signals compared to WT group, indicating a suppression of Ccr modification.

Given that the conversion from Acac to crotonyl intermediates likely involves enzymatic steps, we further investigated the role of key metabolic enzymes. 3-Hydroxybutyrate dehydrogenase 1 (BDH1)^27^ is known to catalyze the reduction of Acac to Bhb, while Enoyl-CoA Hydratase, Short Chain 1 (ECHS1)^28^ is implicated in the subsequent dehydration step to form the crotonate intermediate (Fig 5A). To evaluate their involvement, we generated BDH1 and ECHS1 knockout (KO) cell lines and performed metabolic labeling using Acac-alkyne (Fig 5C). Notably, both KO cell lines exhibited reduced fluorescence signals compared to WT control, indicating a suppression of Ccr modification formation. Together, these findings support a model in which Acac is converted to Bhb and then to crotonyl intermediate that undergoes Michael addition with cysteine residues to form Ccr, with BDH1 and ECHS1 serving as critical enzymatic regulators of this pathway.

## Discussion

Ketone bodies are increasingly produced by the liver during periods of low carbohydrate availability (e.g. fasting, exercise, or ketogenic diets). In addition to serving as energy substrates, ketone bodies function as metabolic regulators that support brain activity, suppress inflammation, modulate immune responses, fuel the heart efficiently and reshape cancer metabolism.^29-33^ Recent studies have identified Kbhb and Kacac as PTMs mediated by ketone bodies, linking them to diverse physiological and pathological processes. In our study, we discovered that ketone bodies can also induce previously uncharacterized cysteine modifications, namely Cys427 and Cys267.

We structurally confirmed the Cys267 modification through both probe-based labeling and synthetic peptide co-elution assays, revealing its origin from the metabolic conversion of Acac into a crotonyl intermediate. Interestingly, a recent study by the Wang group reported carboxyalkylation modifications derived from long-chain fatty acid metabolism using the IsoSTAR platform.^34^ While the modification is related, the cysteine crotonation reported here appears to be distinct in both its metabolic precursor and biosynthetic pathway. Acac is derived from ketogenesis and mainly produced in human liver and then circulate to extrahepatic tissues. We showed Acac is metabolized to Bhb first and subsequently to crotonyl intermediates that covalently react with cysteine thiols to generate Ccr, with BDH1 and ECHS1 as key enzymes for this transformation. These findings identify ketone bodies as novel inducers of cysteine modifications, expanding the known scope of their biochemical reactivity.

Despite these advances, several aspects of the study require further investigation. The precise structure of the Cys427 modification remains unresolved and will necessitate chemical synthesis and structural elucidation. Moreover, although CRISPR-mediated knockout of ECHS1 and BDH1 significantly reduced labeling, in-gel fluorescence signals were still observed, suggesting that compensatory metabolic pathways may also contribute to Ccr formation that need to be further investigated.

Our findings reveal that ketone metabolism can drive novel cysteine modifications that may fundamentally reshape our understanding of ketogenesis and its regulatory functions in biology. Ketone bodies and their downstream metabolites possess chemically reactive functional groups, such as ketones/aldehydes, α,β-unsaturated carboxylic acids, and α-hydrogens adjacent to carbonyls, that can engage in diverse covalent interactions with proteins. These functionalities offer multiple avenues for PTM, either through enzymatic transfer or nonenzymatic reactivity. While lysine acylations such as Kbhb and Kacac have been the primary focus, our findings suggest that other nucleophilic residues, including cysteine, may also serve as modification sites under ketogenic conditions. Given the diversity of ketone-derived metabolites and their metabolic flux in cells, it is likely that additional, currently unrecognized PTMs exist. These findings highlight the need to systematically explore the PTM landscape driven by ketone metabolism and expand our chemical understanding of metabolite-protein interactions.

Interestingly, the Ccr modification identified in this study was exclusively localized to mitochondrial proteins, many of which are enzymes involved in redox balance and energy metabolism. This compartmental specificity implies that the mitochondrial microenvironment, characterized by high thiol content, reactive oxygen species (ROS), and dynamic metabolic flux, may uniquely favor cysteine modification by ketone-derived intermediates. Enzymes such as PRDX3, ACAT1, and HSD17B10, central to detoxification, ketone utilization, and β-oxidation,^35-37^ were among the Ccr-modified targets, suggesting functional relevance to mitochondrial redox regulation. The spatial restriction of this modification raises intriguing questions about its impact on mitochondrial signaling, stress response, and the integration of metabolic and redox control.

## Acknowledgements

We would like to thank Emanuel Petricoin and Weidong Zhou from George Mason University for proteomics analysis. We also want to thank Yi Zheng for illustrations (www.yizhengillustration.com) and Evan R. Roberts and Zhiqun Cindy Zhou for technical support. This research was supported by startup funds from Desai Sethi Urology Institute & Sylvester Comprehensive Cancer Center, University of Miami to Zhipeng A. Wang. Chemical proteomics was supported by startup funds from the University of Florida to Xin Wang.

## Conflicts of interest

There are no conflicts to declare.

---

**Fig S1.**
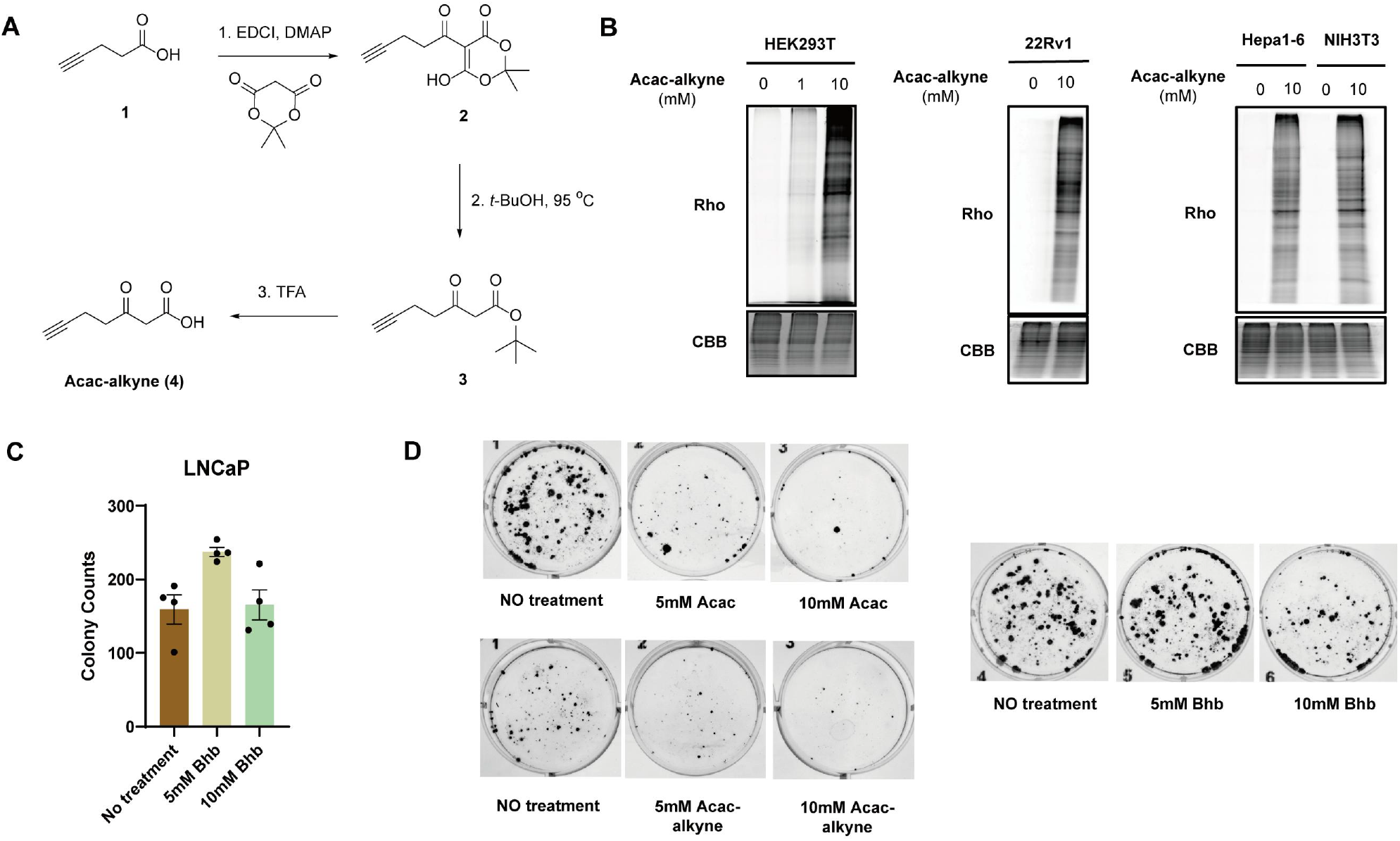
(A) Chemical synthesis of Acac-alkyne. (B) In-gel fluorescence of Acac-alkyne probe in HEK293T, 22Rv1, Hepa1-6 and NIH3T3 cell lines. (C) Bhb do not show colony-suppressive effects. (D) Representative wells from the colony formation assay.

**Fig S2.**
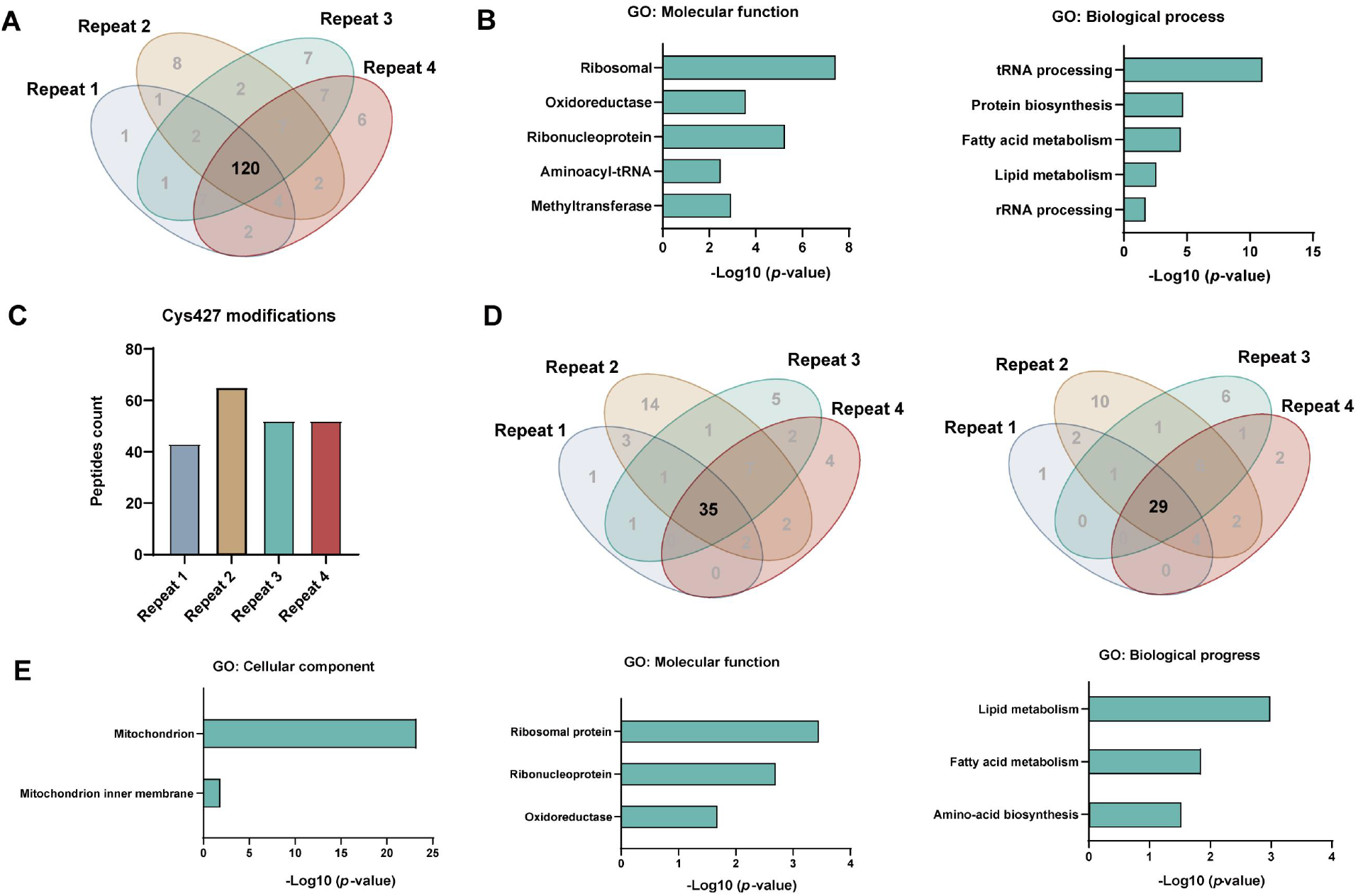
(A) The chemical proteomics data indicated 120 distinct proteins with Cys267 modification across four replicates. (B) GO enrichment analysis of Cys267-modified proteins in the categories of molecular function and biological process. (C, D) The chemical proteomics data showed 35 peptides containing Cys427 modification, mapping to 29 proteins. (E) GO enrichment analysis of Cys427-modified proteins in the categories of cellular component, molecular function and biological process.

**Fig S3.**
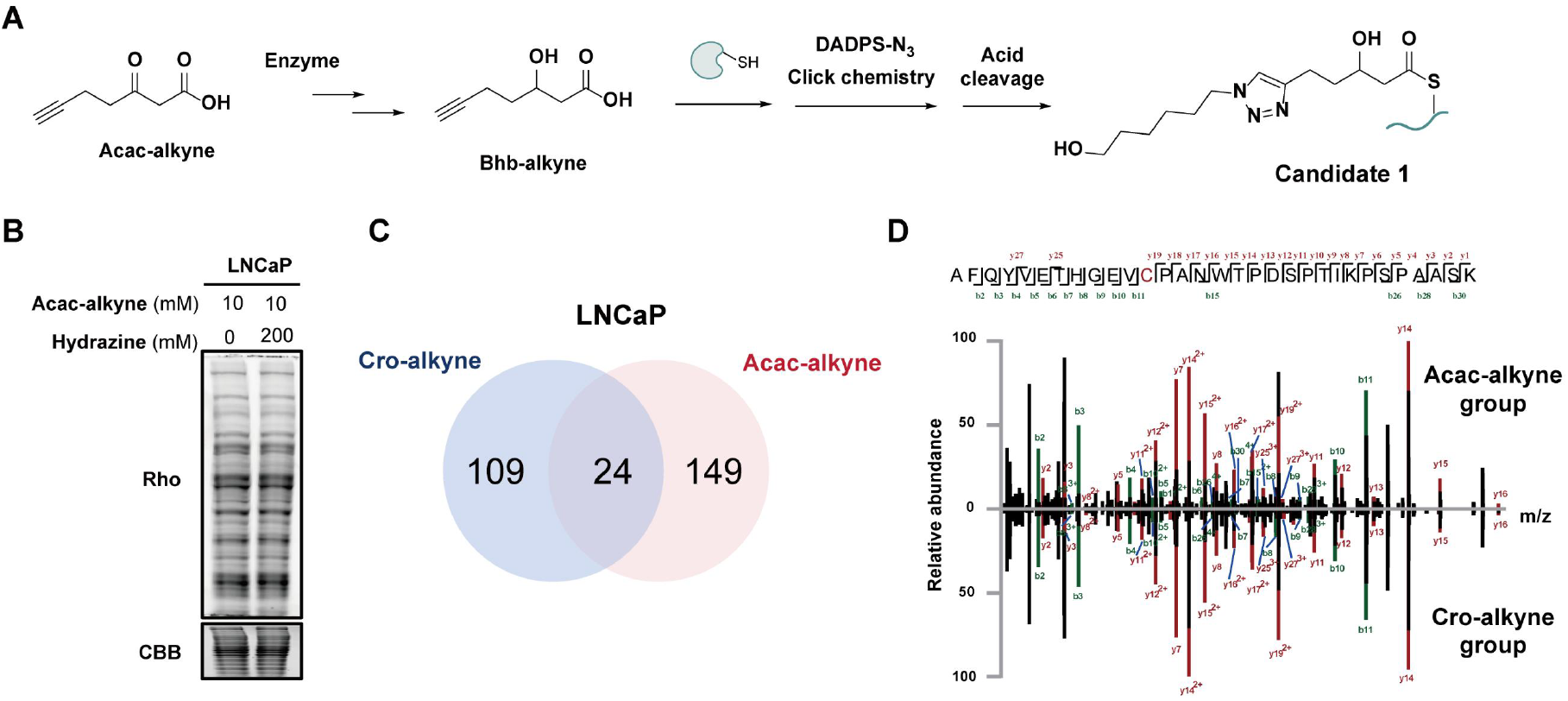
(A) Proposed candidate 1 to form Cys267 modification. (B) In-gel samples treated with 0.2 M hydrazine showed no detectable change in fluorescence intensity. (C) 133 sites were identified to have Cys267 modification in Cro-alkyne group, sharing 24 common sites with that of Acac-alkyne. (D) MS2 spectra of the Cys267 modification at PRDX3_C229 (AFQYVETHGEV**C**PANWTPDSPTIKPSPAASK).

**Fig S4.**
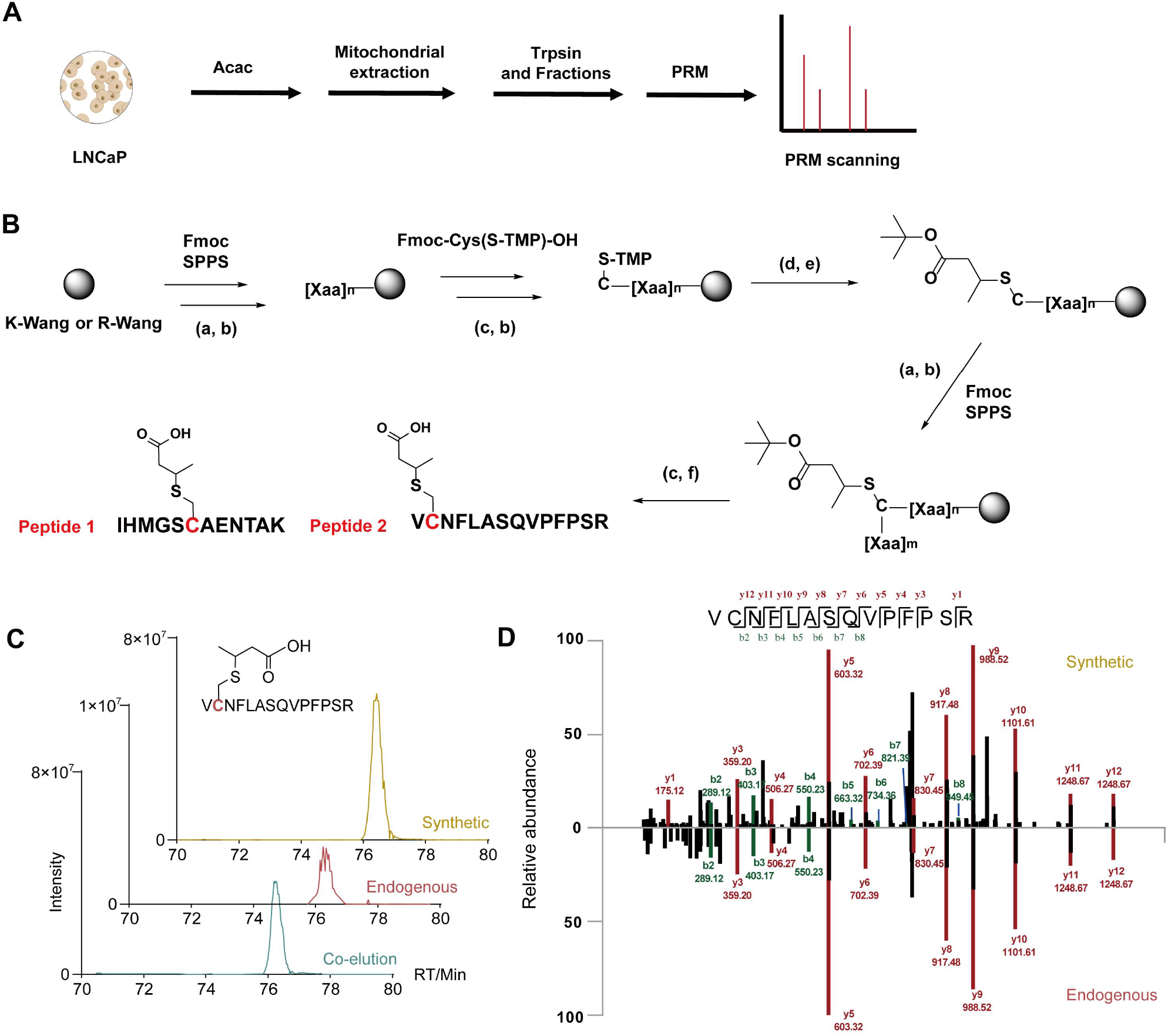
(A) Mitochondrial extracts were analyzed using PRM with an inclusion list incorporating modification site information. (B) Detailed procedures for the synthesis of the two modified peptides. (C) The extracted ion chromatogram corresponding to Ccr modification of HSD17B10_C112 (V**Ccr**NFLASQVPFPSR). (D) MS2 spectra of the Ccr modification at HSD17B10_C112 (V**Ccr**NFLASQVPFPSR).

**Fig S5.**
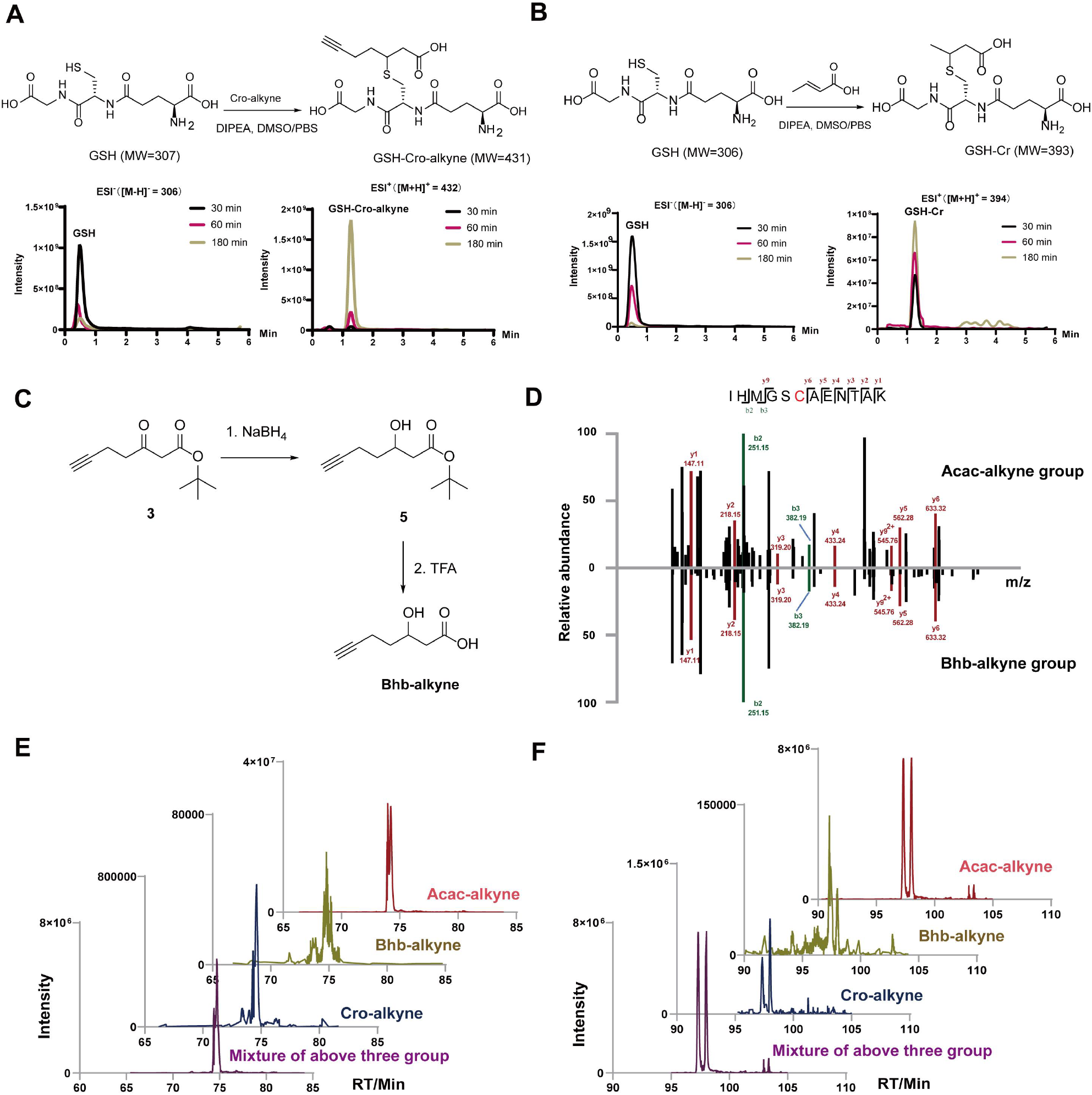
Monitoring the non-enzymatic Michael addition reaction between cro-alkyne (A) or crotonate (B) and GSH in different time points. (C) Synthesis of Bhb-alkyne. (D) MS2 spectra of the probe modification at ACAT1_C126 (IHMGS**Ccr**AENTAK). (E, F) The extracted ion chromatogram corresponding to PRDX3_C229 and HSD17B10_C112 in Acac-alkyne, Bhb-alkyne, Cro-alkyne and mixture groups, respectively.

## Notes

### Competing Interest Statement

The authors have declared no competing interest.

